# SARS-CoV-2 invades host cells via a novel route: CD147-spike protein

**DOI:** 10.1101/2020.03.14.988345

**Authors:** Ke Wang, Wei Chen, Yu-Sen Zhou, Jian-Qi Lian, Zheng Zhang, Peng Du, Li Gong, Yang Zhang, Hong-Yong Cui, Jie-Jie Geng, Bin Wang, Xiu-Xuan Sun, Chun-Fu Wang, Xu Yang, Peng Lin, Yong-Qiang Deng, Ding Wei, Xiang-Min Yang, Yu-Meng Zhu, Kui Zhang, Zhao-Hui Zheng, Jin-Lin Miao, Ting Guo, Ying Shi, Jun Zhang, Ling Fu, Qing-Yi Wang, Huijie Bian, Ping Zhu, Zhi-Nan Chen

## Abstract

Currently, COVID-19 caused by severe acute respiratory syndrome coronavirus 2 (SARS-CoV-2) has been widely spread around the world; nevertheless, so far there exist no specific antiviral drugs for treatment of the disease, which poses great challenge to control and contain the virus. Here, we reported a research finding that SARS-CoV-2 invaded host cells via a novel route of CD147-spike protein (SP). SP bound to CD147, a receptor on the host cells, thereby mediating the viral invasion. Our further research confirmed this finding. First, in vitro antiviral tests indicated Meplazumab, an anti-CD147 humanized antibody, significantly inhibited the viruses from invading host cells, with an EC_50_ of 24.86 μg/mL and IC_50_ of 15.16 μg/mL. Second, we validated the interaction between CD147 and SP, with an affinity constant of 1.85×10^−7^M. Co-Immunoprecipitation and ELISA also confirmed the binding of the two proteins. Finally, the localization of CD147 and SP was observed in SARS-CoV-2 infected Vero E6 cells by immuno-electron microscope. Therefore, the discovery of the new route CD147-SP for SARS-CoV-2 invading host cells provides a critical target for development of specific antiviral drugs.

## INTRODUCTION

In December, 2019, the novel coronavirus (CoV), SARS-CoV-2, has emerged in China [1]. The disease caused by SARS-CoV-2 was named as COVID-19 by the World Health Organization (WHO). As of 10^th^ March 2020, data from the WHO shows there has been an outbreak of COVID-19 in 110 countries and areas around the world, with 113702 confirmed cases. So far there is no specific drug for the treatment of COVID-19. Therefore, intensive researches are urgently needed to elucidate the invasive mechanisms of SARS-CoV-2, thereby providing a potential target for developing specific antiviral drugs.

It has been reported that CoVs are non-segmented positive sense RNA viruses, which primarily cause enzootic infections in birds and mammals and have demonstrated strong lethality in humans [1,2]. These viruses are known to have four structural proteins, including E, M, N and S protein [3,4]. The primary determinant of CoVs tropism is the S protein, which binds to the membrane receptor on the host cells, mediating the viral and cellular membrane fusion [5]. Angiotensin-converting enzyme 2 (ACE2), a homologue of ACE, is one of the important receptors on the cell membrane of the host cells. The interaction of SP and ACE2 contributes to the SARS-CoV invasion for host cells [6]. Interestingly, the structure of SARS-CoV-2 SP is highly similar to that of SARS-CoV SP, and SARS-CoV-2 SP binds to ACE2 with a higher affinity than SARS-CoV SP, indicative of a stronger ability of SAR-CoV-2 to invade host cells [7,8]. In addition to ACE2, other receptors for SP binding on the host cell membrane remain unclear.

CD147, also known as Basigin or EMMPRIN, is a transmembrane glycoprotein that belongs to the immunoglobulin superfamily [9], which is involved in tumor development, plasmodium invasion and virus infection [10-16]. Our previous researches show that CD147 plays a functional role in facilitating SARS-CoV invasion for host cells, and CD147-antagonistic peptide-9 has a high binding rate to HEK293 cells and an inhibitory effect on SARS-CoV [16]. These researches affirm the importance of CD147 in virus invasion for host cells. Owing to the similar characteristics of SARS-CoV and SARS-CoV-2, we conduct the present study to investigate the possible function of CD147 in invasion for host cells by SARS-CoV-2.

In our study, we reported SARS-CoV-2 invaded host cells via a novel route of CD147-SP. Meplazumab, a humanized anti-CD147 antibody, could competitively inhibit the binding of SP and CD147 and prevent the viruses from invading host cells, which provides a potential target for developing antiviral drugs.

## RESULTS

### Blocking CD147 inhibits SARS-CoV-2 replication

The antiviral tests were performed to investigate the possible function of CD147 in SARS-CoV-2 invasion for host cells. Vero E6 cells were cultured in a 96-well plate, and CD147 on Vero E6 cells was blocked with Meplazumab. Then, the cells were infected with SARS-CoV-2 (100TCID_50_) and the cell damage of Vero E6 and viral load were detected by crystal violet staining and quantitative PCR, respectively. Compared with the control group, the inhibitory rate for viruses was significantly increased in CD147-blocking group in a dose-dependent manner, with a concentration for 50% of maximal effect (EC_50_) of 24.86 μg/mL (Figure 1A) and the half maximal inhibitory concentration (IC_50_) of 15.16 μg/mL (Figure 1B). These results show that blocking CD147 on the host cells has an inhibitory effect on SARS-CoV-2, suggesting an essential role of CD147 in facilitating SARS-CoV-2 invasion for host cells.

**Figure 1.**
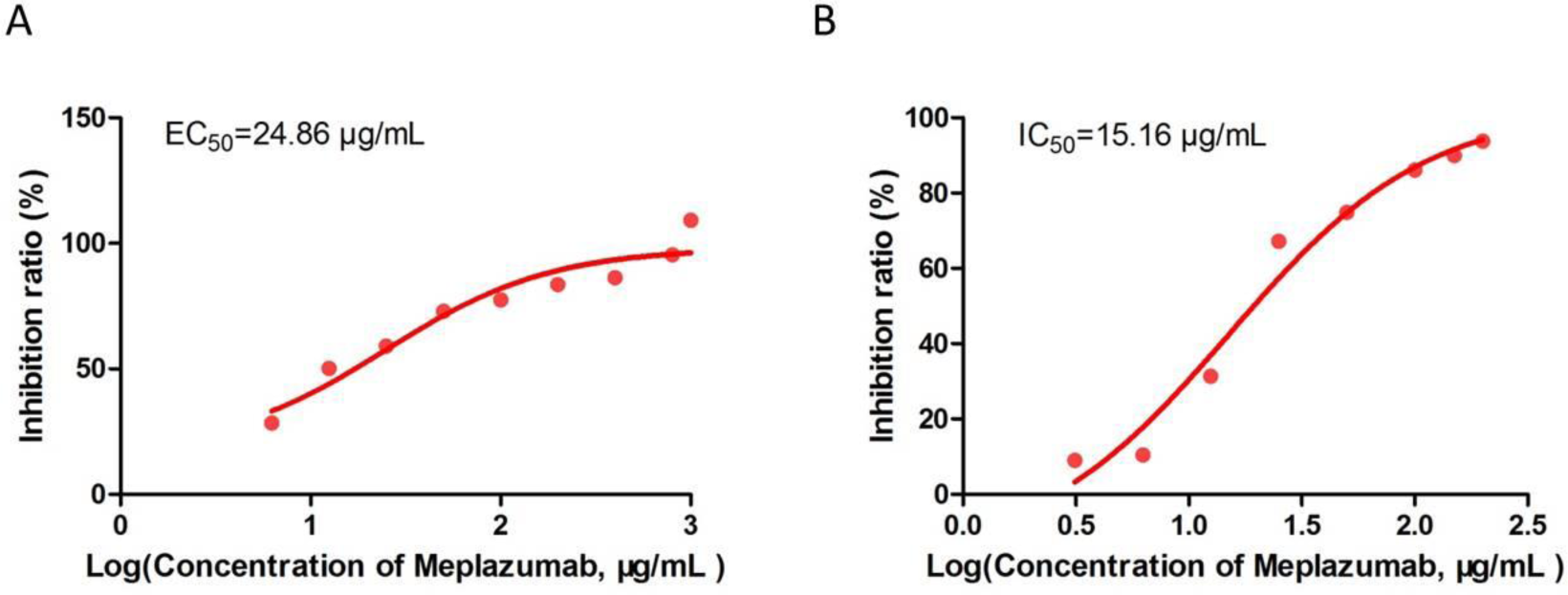
Meplazumab inhibits the SARS-CoV-2 replication. **(A and B)** 1×10^4^ Vero E6 cells were cultured in a 96-well plate at 37°C overnight, the supernatant was discard and 100 μl of medium (containing different concentrations of Meplazumab) was added into the plates to incubate for 1 h. Then the cells were infected with SARS-CoV-2 (100TCID_50_). After one-hour infection, the supernatants were removed and 200 μl of medium (containing different concentrations of Meplazumab) was added, the cells were cultured to observe the cytopathic changes for 2-3 days. The supernatants were harvested to detect the gene copy number of virus with quantitative PCR and the Vero E6 cells were stained by crystal violet staining. Finally, the value of optical density (OD) at 570 nm was measured with a microplate reader.

### Identification of interaction between CD147 and SP

CD147 is closely associated with SARS-CoV-2 invasion for host cells, and SP binds to cellular receptors to mediate infection of host cells [7], leading to the question whether there is some relationship between CD147 and SP that affects the invasion of the SARS-CoV-2. Therefore, the interaction assays was conducted to clarify the interaction of the two proteins. Surface Plasmon Resonance (SPR) assay confirmed the interaction of CD147 and SP (RBD), with an affinity constant of 1.85×10^−7^M (Figure 2A). In Co-IP assay, anti-SP antibody and anti-CD147 antibody were used for antibody immobilization, respectively. The mIgG and rIgG were selected as negative control. Then, the mixture of His-CD147 and His-SP (RBD) was added in each group for IP, and the eluted protein was detected by western blot. The result showed that there was an interaction between CD147 and SP (RBD), which was consistent with the SPR assay (Figure 2B). In addition, the interaction of CD147-SP (RBD) was also confirmed by ELISA assay (Figure 2C). In the meanwhile, competitive inhibition assay showed that Meplazumab could competitively inhibit the binding of SP and CD147, with an IC_50_ of 16.44 μg/mL (Figure 2D). These results confirm the interaction of CD147 and SP, indicating a novel potential route for SARS-CoV-2 to invade host cells.

**Figure 2.**
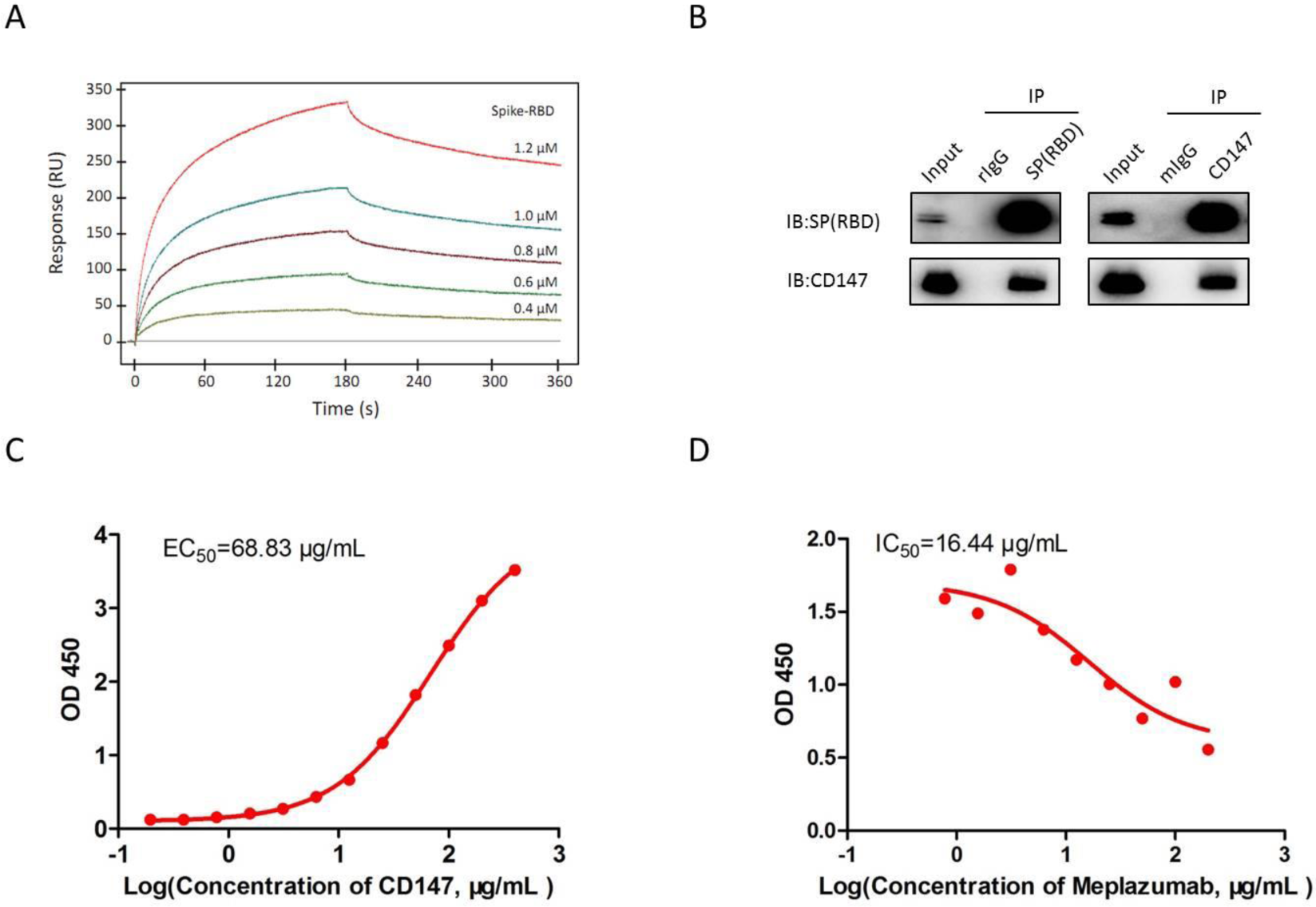
Identification of interaction between CD147 and SP. **(A)** The interaction of CD147 and SP detected by SPR assay, KD = 1.85×10^−7^M. **(B)** The interaction of CD147 and SP detected by Co-IP assay. Anti-CD147 antibody and anti-SARS-CoV-2 Spike antibody were used for antibody immobilization for Co-IP. The mIgG and rIgG were selected as negative control. **(C)** The interaction of CD147 and SP detected by ELISA. **(D)** The ability of Meplazumab to compete with SP for CD147 binding performed by competitive inhibition ELISA.

### Detection of localization of CD147 and SP by immuno-electron microscope

To further determine the localization of CD147 and SP in the process of viral infection, the Vero E6 cells infected with SARS-CoV-2 were observed by immune-electron microscope. After traced by 10 nm (CD147) and 20 nm (SP) gold colloid-labeled antibodies, we found that the two proteins mainly presented in viral inclusion bodies of Vero E6 cells (Figure 3A-C). These results reinforce the finding that the CD147-SP interaction enhances viral invasion for host cells.

**Figure 3.**
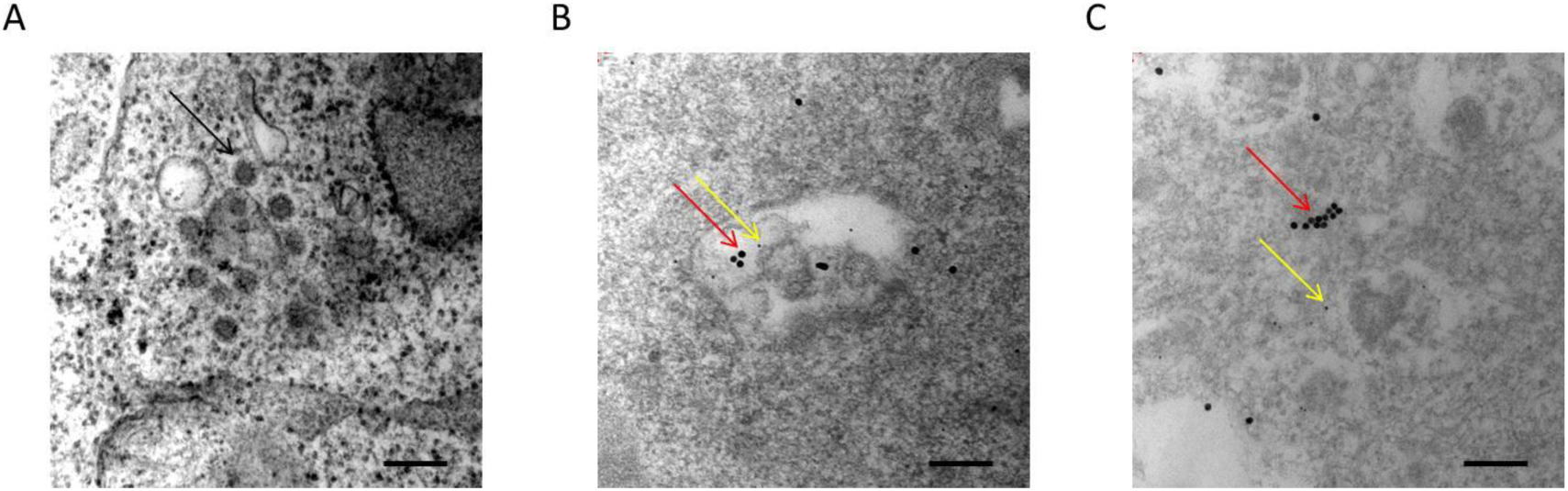
The localization of CD147 and SP observed by immune-electron microscope. **(A)** The SARS-CoV-2 infected Vero E6 cell with visible SARS-CoV-2 virion (black arrow). (**B** and **C**) The localization of CD147 (10nm-gold colloid, yellow arrows) and SP (20nm-gold colloid, red arrows) in viral inclusion bodies of SARS-CoV-2 infected Vero E6 cell. Scale bar = 200nm.

## DISCUSSION

The emergence of SARS-CoV-2 and its rapid spread across the world has triggered a global health emergency. Patients with COVID-19 show abnormal findings on chest computed tomography, along with common symptoms that include cough, fever and fatigue [17]. Middle-aged and elderly patients with underlying comorbidities are susceptible to respiratory failure and may have a poorer prognosis [18]. The pathological features of patients with COVID-19 are very similar to those of patients with SARS and MERS infection [19]. However, COVID-19 is more contagious than SARS and MERS [20]. Therefore, it is urgent to develop a specific antiviral drug against COVID-19.

It is known that ACE2 has been widely recognized as an essential receptor in virus invasion, and viral replication was specifically inhibited by an anti-ACE2 antibody [21]. However, ACE2 is widely distributed in variety of tissues, especially in the heart, kidney and testes [22], which plays a profound role in controlling blood pressure, preventing heart failure and kidney injury [23-25]. For lung diseases, the loss of ACE2 enhances vascular permeability and lung edema, activates the renin-angiotensin system and contributes to pathogenesis of severe lung injury [26]. Therefore, treatment with ACE2 as target may have a negative effect on its protective role.

In our study, CD147 was identified as a novel receptor of SP, and the CD147-SP interaction facilitated the viral invasion for host cells. As mentioned earlier, CD147 is highly expressed in tumor tissues, inflamed tissues and pathogen infected cells, which leads to low cross-reaction with normal cells [27,28]. In the meanwhile, Meplazumab, a humanized anti-CD147 antibody, could effectively inhibit the viruses from invading host cells by blocking CD147. Our previous research and development of original antibody drugs, such as Metuximab, Metuzumab, Meplazumab, all exhibits sound safety in preclinical research and clinical administration. Therefore, the CD147 targeted drug is safe and reliable in its application, and is a receptor-blocking drug, which blocks the virus from invading host cells, without being affected by virus variation.

In general, our study reported that CD147 was a novel receptor for SP. The route of CD147-SP facilitated SARS-CoV-2 invasion for host cells. Therefore, this route provides a novel target, with a good druggability, for specific antiviral drug development in the treatment of COVID-19.

## METHODS

### Ethics statement

This study was approved by the ethics committee of First Affiliated Hospital of Fourth Military Medical University (KY20202005-1). SARS-CoV-2 strain (2019-nCoV BetaCoV /Beijing /AMMS01 /2020) was isolated and preserved in State Key Laboratory of Pathogen and Biosecurity at Beijing Institute of Microbiology and Epidemiology. All the experiments involved with viruses were performed in BSL-3 Laboratory of Beijing Institute of Microbiology and Epidemiology.

### Cell culture

The cell lines (Vero E6 and 293T) have been tested and authenticated using Short Tandem Repeat DNA profiling by Beijing Microread Genetics Co., Ltd (Beijing, China) and were cultured at 37°C under 5% CO_2_ in DMEM or RPMI 1640 medium supplemented with 10% fetal bovine serum, 1% penicillin/streptomycin and 2% L-glutamine.

### In vitro antiviral tests

1×10^4^ Vero E6 cells were cultured in a 96-well plate at 37°C overnight, the supernatant was discard and 100 μl of medium (containing 3.125, 6.25, 12.5, 25, 50, 100, 150, 200 μg/mL Meplazumab) was added into the plates to incubate for 1 h. Then the cells were infected with SARS-CoV-2 (100TCID_50_). After one-hour infection, the supernatants were removed and 200 μl of medium (containing 3.125, 6.25, 12.5, 25, 50, 100, 150, 200 μg/mL Meplazumab) was added, the cells were cultured to observe the cytopathic changes for 2-3 days. The supernatants were harvested to detect the gene copy number of virus with quantitative PCR. Finally, the value of optical density (OD) at 570 nm was measured with a microplate reader (Epoch, BioTek Instruments, Inc.). A virus control group and a blank control group were set at the same time. The same method was conducted with different antibody concentration (containing 6.25, 12.5, 25, 50, 100, 200, 400, 800, 1000 μg/mL Meplazumab), and the Vero E6 cells were stained by crystal violet staining.

### SPR assay

SPR analysis was performed by BIAcore 3000 system (BIAcore). His-CD147 (produced by our laboratory) was fixed to the surface of CM5 sensor chips (GE Healthcare Bio-Sciences AB) by amino coupling kit (GE Healthcare, BR-1000-50); interaction between CD147 and SP (RBD) (GenScript, China) was detected using Kinetic Analysis/Concentration Series/Direct Binding mode, and the flow rate was set to 15 μl/min, both binding time and dissociation time were 3 min. The results were analyzed by BIAevaluation software to determine the affinity constant.

### Co-IP assay

Co-IP assay was performed using Pierce^®^ Co-Immunoprecipitation Kit (26149, Thermo Fisher Scientific, USA) according to the manufacturer’s protocol. Anti-CD147 antibody (HAb18, produced by our laboratory) and anti-SARS-CoV-2 Spike antibody (40150-R007, Sino Biological, China) were used for antibody immobilization for Co-IP. The eluted proteins were detected by western blot. After boiling for 5-10 minutes, the eluted proteins were loaded to 12% SDS-PAGE gel and then transferred to PVDF membranes (Millipore, Boston, USA). After blocking with 5% non-fat milk for 1 hour, the membrane was incubated with the corresponding primary antibodies (anti-CD147, HAb18, produced by our laboratory, dilution 1:2000; anti-SARS-CoV-2 Spike antibody, 40150-R007, Sino Biological, China, dilution 1:2000) at 4°C overnight. The images were developed after incubation with the secondary antibodies (goat anti-mouse IgG(H+L) antibody, 31430, Thermo Fisher Scientific, MA, USA, dilution 1:5000; goat anti-rabbit IgG(H+L) antibody; 31460, Thermo Fisher Scientific, MA, USA, dilution 1:5000) at room temperature for 1 hour.

### ELISA

ELISA assay was performed to identify the interaction of CD147 and SP. The His-SP (RBD) protein was coated on microplate, and then incubated with different concentrations of His-CD147 protein (2-fold dilution, 400-0.195 μg/mL) at 37°C for 1 hour. After washing with PBST, the samples were incubated with anti-CD147 antibody (HAb18, produced by our laboratory) for 1 hour; and then incubated with horseradish peroxidase (HRP)-labeled goat anti-mouse antibody for 1 hour. After coloration, the OD value at 450 nm was measured with a full-wavelength microplate reader (Epoch, BioTek Instruments, Inc.). The ability of Meplazumab to compete with SP for CD147 binding was performed by competitive inhibition ELISA. The protein His-CD147 was coated onto the wells of microplate, the mixture of His-SP (RBD) and Meplazumab (2-fold dilution, 200-0.781 µg/mL) was added and incubated. Subsequently, the samples were incubated with anti-SARS-CoV-2 Spike antibody (40150-R007, Sino Biological, China) and then detected with HRP-labeled goat anti-rabbit antibody for 1 hour at 37°C, the following steps were similar to those described earlier.

### Immuno-electron microscope

Vero E6 cells infected with SARS-CoV-2 were harvested and treated with fixative at 4°C. The sections of the cells were prepared through dehydration and embedding. After washing with ultrapure water, the sections were treated with 1% H_2_O_2_ for 10 min. Having been blocked with goat serum for 30min, the sections was incubated with corresponding primary antibodies (anti-CD147, HAb18, produced by our laboratory, dilution 1:200; anti-SARS-CoV-2 Spike antibody, 40150-R007, Sino Biological, China, dilution 1:400) for 16 hours at 4°C overnight. Subsequently, the sections were washed with PBS for 5 min, and treated with PBS (containing 1% bovine serum albumin, BSA, pH 8.2) for 7 min. Then the gold colloid-labeling method was used to determine the localization of CD147 and SP for 1h at room temperature, and the sections were consecutively stained with 5% uranium acetate and lead acetate. Immune-electron microscope was used to observe the protein location.

### Statistical Analysis

All the statistical analyses were performed using GraphPad prism 5.0. Data from ELISA were analyzed using parameter curve fitting.

## ACKNOWLEDGEMENTS

The authors thank Prof. Nanshan Zhong and members of Zhong lab for their critical guidance.

## REFERENCES

1. Huang C, Wang Y, Li X, Ren L, Zhao J, Hu Y, Zhang L, Fan G, Xu J, Gu X, Cheng Z, Yu T, Xia J, et al. Clinical features of patients infected with 2019 novel coronavirus in Wuhan, China. LANCET. 2020; 395: 497–506.

2. Schoeman D, Fielding BC. Coronavirus envelope protein: current knowledge. VIROL J. 2019; 16: 69.

3. Rota PA, Oberste MS, Monroe SS, Nix WA, Campagnoli R, Icenogle JP, Penaranda S, Bankamp B, Maher K, Chen MH, Tong S, Tamin A, Lowe L, et al. Characterization of a novel coronavirus associated with severe acute respiratory syndrome. SCIENCE. 2003; 300: 1394–1399.

4. Thiel V, Ivanov KA, Putics A, Hertzig T, Schelle B, Bayer S, Weissbrich B, Snijder EJ, Rabenau H, Doerr HW, Gorbalenya AE, Ziebuhr J. Mechanisms and enzymes involved in SARS coronavirus genome expression. J GEN VIROL. 2003; 84: 2305–2315.

5. Hulswit RJ, de Haan CA, Bosch BJ. Coronavirus Spike Protein and Tropism Changes. ADV VIRUS RES. 2016; 96: 29–57.

6. Hui DS. Epidemic and Emerging Coronaviruses (Severe Acute Respiratory Syndrome and Middle East Respiratory Syndrome). CLIN CHEST MED. 2017; 38: 71–86.

7. Wrapp D, Wang N, Corbett KS, Goldsmith JA, Hsieh CL, Abiona O, Graham BS, McLellan JS. Cryo-EM structure of the 2019-nCoV spike in the prefusion conformation. SCIENCE. 2020.

8. Yan R, Zhang Y, Li Y, Xia L, Guo Y, Zhou Q. Structural basis for the recognition of the SARS-CoV-2 by full-length human ACE2. SCIENCE. 2020.

9. Cui J, Huang W, Wu B, Jin J, Jing L, Shi WP, Liu ZY, Yuan L, Luo D, Li L, Chen ZN, Jiang JL. N-glycosylation by N-acetylglucosaminyltransferase V enhances the interaction of CD147/basigin with integrin beta1 and promotes HCC metastasis. J PATHOL. 2018; 245: 41–52.

10. Huang Q, Li J, Xing J, Li W, Li H, Ke X, Zhang J, Ren T, Shang Y, Yang H, Jiang J, Chen Z. CD147 promotes reprogramming of glucose metabolism and cell proliferation in HCC cells by inhibiting the p53-dependent signaling pathway. J HEPATOL. 2014; 61: 859–866.

11. Zhao P, Zhang W, Wang SJ, Yu XL, Tang J, Huang W, Li Y, Cui HY, Guo YS, Tavernier J, Zhang SH, Jiang JL, Chen ZN. HAb18G/CD147 promotes cell motility by regulating annexin II-activated RhoA and Rac1 signaling pathways in hepatocellular carcinoma cells. HEPATOLOGY. 2011; 54: 2012–2024.

12. Lu M, Wu J, Hao ZW, Shang YK, Xu J, Nan G, Li X, Chen ZN, Bian H. Basolateral CD147 induces hepatocyte polarity loss by E-cadherin ubiquitination and degradation in hepatocellular carcinoma progress. HEPATOLOGY. 2018; 68: 317–332.

13. Zhang MY, Zhang Y, Wu XD, Zhang K, Lin P, Bian HJ, Qin MM, Huang W, Wei D, Zhang Z, Wu J, Chen R, Feng F, et al. Disrupting CD147-RAP2 interaction abrogates erythrocyte invasion by Plasmodium falciparum. BLOOD. 2018; 131: 1111–1121.

14. Pushkarsky T, Zybarth G, Dubrovsky L, Yurchenko V, Tang H, Guo H, Toole B, Sherry B, Bukrinsky M. CD147 facilitates HIV-1 infection by interacting with virus-associated cyclophilin Proc Natl Acad Sci U S A. 2001; 98: 6360–6365.

15. Castro AP, Carvalho TM, Moussatche N, Damaso CR. Redistribution of cyclophilin A to viral factories during vaccinia virus infection and its incorporation into mature particles. J VIROL. 2003; 77: 9052–9068.

16. Chen Z, Mi L, Xu J, Yu J, Wang X, Jiang J, Xing J, Shang P, Qian A, Li Y, Shaw PX, Wang J, Duan S, et al. Function of HAb18G/CD147 in invasion of host cells by severe acute respiratory syndrome coronavirus. J INFECT DIS. 2005; 191: 755–760.

17. Jiang S, Xia S, Ying T, Lu L. A novel coronavirus (2019-nCoV) causing pneumonia-associated respiratory syndrome. CELL MOL IMMUNOL. 2020.

18. Liu K, Fang YY, Deng Y, Liu W, Wang MF, Ma JP, Xiao W, Wang YN, Zhong MH, Li CH, Li GC, Liu HG. Clinical characteristics of novel coronavirus cases in tertiary hospitals in Hubei Province. Chin Med J (Engl). 2020.

19. Xu Z, Shi L, Wang Y, Zhang J, Huang L, Zhang C, Liu S, Zhao P, Liu H, Zhu L, Tai Y, Bai C, Gao T, et al. Pathological findings of COVID-19 associated with acute respiratory distress syndrome. Lancet Respir Med. 2020.

20. [The epidemiological characteristics of an outbreak of 2019 novel coronavirus diseases (COVID-19) in China]. Zhonghua Liu Xing Bing Xue Za Zhi. 2020; 41: 145–151.

21. Li W, Moore MJ, Vasilieva N, Sui J, Wong SK, Berne MA, Somasundaran M, Sullivan JL, Luzuriaga K, Greenough TC, Choe H, Farzan M. Angiotensin-converting enzyme 2 is a functional receptor for the SARS coronavirus. NATURE. 2003; 426: 450–454.

22. Tipnis SR, Hooper NM, Hyde R, Karran E, Christie G, Turner AJ. A human homolog of angiotensin-converting enzyme. Cloning and functional expression as a captopril-insensitive carboxypeptidase. J BIOL CHEM. 2000; 275: 33238–33243.

23. Wong DW, Oudit GY, Reich H, Kassiri Z, Zhou J, Liu QC, Backx PH, Penninger JM, Herzenberg AM, Scholey JW. Loss of angiotensin-converting enzyme-2 (Ace2) accelerates diabetic kidney injury. AM J PATHOL. 2007; 171: 438–451.

24. Rentzsch B, Todiras M, Iliescu R, Popova E, Campos LA, Oliveira ML, Baltatu OC, Santos RA, Bader M. Transgenic angiotensin-converting enzyme 2 overexpression in vessels of SHRSP rats reduces blood pressure and improves endothelial function. HYPERTENSION. 2008; 52: 967–973.

25. Der Sarkissian S, Grobe JL, Yuan L, Narielwala DR, Walter GA, Katovich MJ, Raizada MK. Cardiac overexpression of angiotensin converting enzyme 2 protects the heart from ischemia-induced pathophysiology. HYPERTENSION. 2008; 51: 712–718.

26. Kuba K, Imai Y, Ohto-Nakanishi T, Penninger JM. Trilogy of ACE2: a peptidase in the renin-angiotensin system, a SARS receptor, and a partner for amino acid transporters. Pharmacol Ther. 2010; 128: 119–128.

27. Kosugi T, Maeda K, Sato W, Maruyama S, Kadomatsu K. CD147 (EMMPRIN/Basigin) in kidney diseases: from an inflammation and immune system viewpoint. Nephrol Dial Transplant. 2015; 30: 1097–1103.

28. Su H, Yang Y. The roles of CyPA and CD147 in cardiac remodelling. EXP MOL PATHOL. 2018; 104: 222–226.

